# Notable sequence homology of the ORF10 protein introspects the architecture of SARS-COV-2

**DOI:** 10.1101/2020.09.06.284976

**Authors:** Sk. Sarif Hassan, Diksha Attrish, Shinjini Ghosh, Pabitra Pal Choudhury, Vladimir N. Uversky, Bruce D. Uhal, Kenneth Lundstrom, Nima Rezaei, Alaa A. A. Aljabali, Murat Seyran, Damiano Pizzol, Parise Adadi, Tarek Mohamed Abd El-Aziz, Antonio Soares, Ramesh Kandimalla, Murtaza Tambuwala, Amos Lal, Gajendra Kumar Azad, Samendra P. Sherchan, Wagner Baetas-da-Cruz, Giorgio Palù, Adam M. Brufsky

## Abstract

The global public health is endangered due to COVID-19 pandemic, which is caused by Severe Acute Respiratory Syndrome Coronavirus-2 (SARS-CoV-2). Despite having similar pathology to MERS and SARS-CoV, the infection fatality rate of SARS-CoV-2 is likely lower than 1%. SARS-CoV-2 has been reported to be uniquely characterized by the accessory protein ORF10, which contains eleven cytotoxic T lymphocyte (CTL) epitopes of nine amino acids length each, across various human leukocyte antigen (HLA) subtypes. In this study, all missense mutations found in sequence databases were examined across twnety-two unique SARS-CoV-2 ORF10 variants that could possibly alter viral pathogenicity. Some of these mutations decrease the stability of ORF10, e.g. I4L and V6I were found in the MoRF region of ORF10 which may also possibly contribute to Intrinsic protein disorder. Furthermore, a physicochemical and structural comparative analysis was carried out on SARS-CoV-2 and Pangolin-CoV ORF10 proteins, which share 97.37% amino acid homology. The high degree of physicochemical and structural similarity of ORF10 proteins of SARS-CoV-2 and Pangolin-CoV open questions about the architecture of SARS-CoV-2 due to the disagreement of these two ORF10 proteins over their sub-structure (loop/coil region), solubility, antigenicity and change from the strand to coil at amino acid position 26, where tyrosine is present. Altogether, SARS-CoV-2 ORF10 is a promising pharmaceutical target and a protein which should be monitored for changes which correlate to change pathogenesis and clinical course of COVID-19 infection.

## Introduction

Severe Acute Respiratory Syndrome Coronavirus-2 (SARS-CoV-2), responsible for the global pandemic, has brought the whole world to a stand-still^1,2^. The contagious nature of this virus is concerning as it has infected more than 25 million people worldwide claiming 850,000 deaths, so far^3–5^. In addition to low-pathogenicity and endemic coronaviruses, high pathogenic Severe Acute Respiratory Syndrome Coronavirus (SARS-CoV) and Middle East Respiratory Syndrome Coronavirus (MERS-CoV) in 2002 and 2013, respectively, caused severe human illnesses, e.g. pneumonia, and renal failure but without any pandemic grade transmission capacities^6^. SARS-CoV had a 9.7% infection fatality rate and MERS a 37% infection fatality rate, but SARS-CoV-2 has a lower than 1% infection fatality^7^. Therefore, it is vital to monitor critical mutations in the proteins such as ORF10 (open reading frame 10) that could possibly change viral pathogenicity. SARS-CoV-2 is a Baltimore class IV positive-sense, single-stranded RNA virus with four structural proteins, sixteen non-structural proteins, and six accessory proteins^8^.

The smallest accessory protein in SARS-CoV-2, the 38-residue peptide ORF10, and distinguishes the infection more rapidly than PCR based strategies^9^. The protein SARS-CoV-2 ORF10 has the highest number of immunogenic epitopes of all putative ORF proteins, therefore making it a potential target for vaccine development^10^. Due to its short length, ORF10 has been suggested to be an insertion mutation. However, this is unlikely as the ORF10 gene is present at the terminal its sgRNA sequence. It has been hypothesised that ORF10 is a transposon, but this is also unlike as transposons are of larger size^9^.

ORF10 consists of a Molecular Recognition Feature (MoRF) region from amino acid residue 3 to 7, which is a molecular recognition site for interaction with other proteins^11^. It is one of the critical properties of intrinsically disordered proteins that allow proteins to adapt an ensemble of conformations when bound to different proteins, and this permits interaction with multiple proteins^12^. Through high-throughput analysis it was revealed that ORF10 can interact with a large number of host proteins despite its small structure; therefore, this aspect can be likely attributed to the MoRF region^11^. Through bioinformatics, it was previously reported that the SARS-CoV-2 ORF10 exhibits interaction with multiple members of the Cullin-ubiquitin-ligase complex and controls the host-ubiquitin machinery for viral pathogenesis^13–16^.

Humans may not have been able to utilize any memory B and T cells elicited against other microorganisms to target ORF10 and fight SARS-CoV-2, contributing to its contagious nature^17^. It was further reported that no sequence homology was found with any protein in the NCBI protein depository. Recently, SARS-CoV-2 ORF10 is found to have 99.15% nucleotide similarity to that of Pangolin-CoV-2020^18,19^.

The present study examines mutations discovered in SARS-CoV-2 ORF10 variants, which along with their physiochemical and immunological properties suggests the significance of these mutations to alter pathogenesis and to possibly identify some potential vaccine candidates. A inclusive parity and disparity analysis between the two ORF10 proteins of SARS-CoV-2 and Pangolin-CoV was also conducted.

## Results

### Mutations in SARS-CoV-2 ORF10

Each unique ORF10 sequence was aligned using the National Center for Biotechnology Information (NCBI) protein p-blast and omega blast suites to determine the mismatches and thereby, the missense mutations (amino acid changes) were identified^20,21^ (Figure 1(A)). A mutation from one amino acid *A*_1_ to another *A*_2_ at the position *p* is denoted by *A*_1_ *pA*_2_ or *A*_1_(*p*)*A*_2_. Based on the mutations, conserved and non-conserved residues in ORF10 proteins are identified and marked in different colors in (Figure 2(B). Also, the molecular recognition features (MoRF) (YINVF) are predicted using the server MoRFchibi for the ORF10 Wuhan sequence^22^.

**Figure 1.**
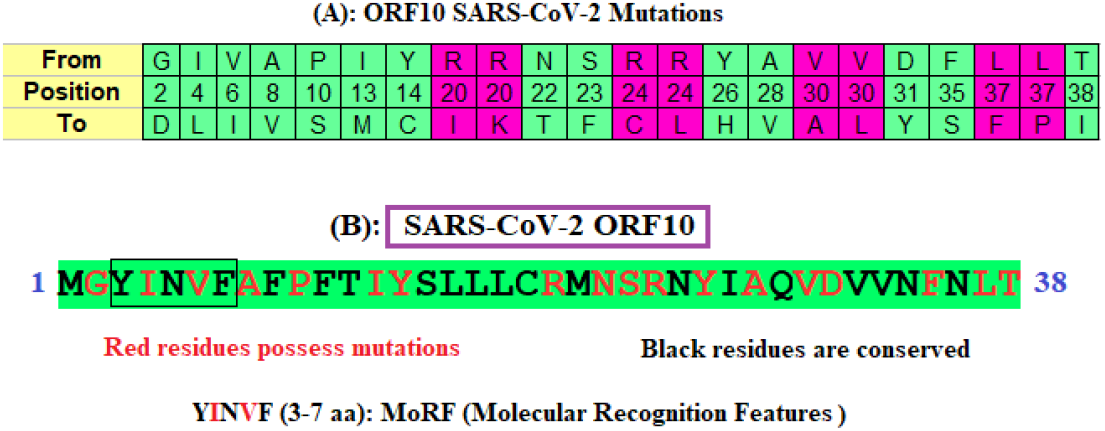
(A): Mutations and their amino acid positions in ORF10 proteins of SARS-CoV-2; (B): Conserved, mutated residues and molecular recognition features of ORF10 (YP_009725255) of SARS-CoV-2.

**Figure 2.**
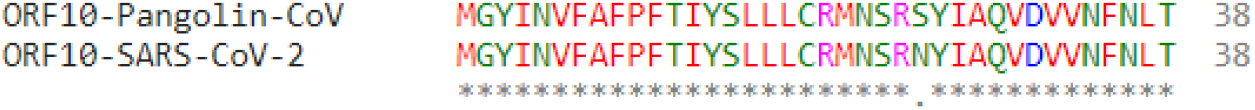
Alignment of two ORF10 sequences (37 out of 38 identical residues) of Pangolin-CoV.

There are altogether 22 distinct missense mutations which were examined across 22 unique ORF10 variants of SARS-CoV-2. These missense mutations are found in the entire ORF10 sequence starting from the amino acid position 2 to 38. The amino acids arginine (R), valine (V), and leucine (L) are substituted to more than one amino acid at fixed positions (marked magenta in Figure 1(B). The largest conserved region across all the 24 ORF10 variants is “SLLLC” at positions 15–19.

Note that each unique variant (Table 2) of SARS-CoV-2 ORF10 possesses a single missense mutation (Table 1).

**Table 1.** Twenty-two ORF10 proteins (SARS-CoV-2) and their corresponding mutations and predicted effects with changes in chemical properties. ^*^PROVEAN score: If the PROVEAN score is equal to or below a predefined threshold (e.g., −2.5), the protein variant is predicted to have a “deleterious” effect. If the PROVEAN score is above the threshold, the variant is predicted to have a “neutral” effect. ^*^RI: Reliability Index ranges from 0 to 9.

| Accession ID | Mutations | Type of Mutations | *PROVEAN Score | Effect of Mutations on Structure | * RI | Polarity Changes | Charge |
| --- | --- | --- | --- | --- | --- | --- | --- |
| QNI23218.1 | G2D | Deleterious | -7 | Decrease | 7 | NP to P | Neutral to Acidic |
| QIS29991.1 | V6I | Neutral | -1 | Decrease | 7 | NP to NP | Neutral to Neutral |
| QLI33453.1 | Y14C | Deleterious | -9 | Decrease | 2 | P to P | Neutral to Neutral |
| QNC04532.1 | R20I | Deleterious | -8 | Decrease | 3 | NP to NP | Basic (strongly) to Neutral |
| QLA48060.1 | R20K | Deleterious | -3 | Decrease | 8 | NP to P | Basic (strongly) to Basic |
| QMT97141.1 | S23F | Deleterious | -6 | Increase | 2 | P to NP | Neutral to Neutral |
| QMU93213.1 | R24C | Deleterious | -8 | Decrease | 7 | P to P | Basic (strongly) to Neutral |
| QMT54534.1 | R24L | Deleterious | -7 | Decrease | 9 | P to NP | Basic (strongly) to Neutral |
| QKU54102.1 | Y26H | Deleterious | -5 | Decrease | 8 | P to P | Neutral to Basic (weakly) |
| QNI25281.1 | V30A | Deleterious | -4 | Decrease | 9 | NP to NP | Neutral to Neutral |
| QNC49349.1 | V30L | Deleterious | -3 | Decrease | 4 | NP to NP | Neutral to Neutral |
| QNA70543.1 | L37F | Deleterious | -4 | Decrease | 7 | NP to NP | Neutral to Neutral |
| QKV37245.1 | T38I | Deleterious | -6 | Decrease | 5 | P to NP | Neutral to Neutral |
| QKV08176.1 | L37P | Deleterious | -7 | Decrease | 8 | NP to NP | Neutral to Neutral |
| QNB17780.1 | F35S | Deleterious | -8 | Decrease | 9 | NP to P | Neutral to Neutral |
| QMT94417.1 | D31Y | Deleterious | -9 | Decrease | 6 | P to P | Acidic to Neutral |
| QLY88596.1 | A28V | Deleterious | -4 | Decrease | 5 | NP to NP | Neutral to Neutral |
| QLG76514.1 | N22T | Deleterious | -6 | Decrease | 1 | P to P | Neutral to Neutral |
| QLG99793.1 | I13M | Deleterious | -3 | Decrease | 8 | NP to NP | Neutral to Neutral |
| QNG42985.1 | P10S | Deleterious | -8 | Decrease | 8 | NP to P | Neutral to Neutral |
| QLJ57416.1 | A8V | Deleterious | -4 | Increase | 3 | NP to NP | Neutral to Neutral |
| QNG41574.1 | I4L | Neutral | -2 | Increase | 1 | NP to NP | Neutral to Neutral |

**Table 2.** Twenty-four unique ORF10 protein IDs with associated geo-location and date of collection of the sample

| Accession | Geo_location | Collection_Date | Accession | Geo_location | Collection_Date |
| --- | --- | --- | --- | --- | --- |
| YP_009725255 | China | 2019-12 | QNC04532 | USA | 2020-04-29 |
| QLJ57416 | USA: WA | 2020 | QNI25281 | USA: Virginia | 2020-05 |
| QIS29991 | China: Hubei, Wuhan | 2020-01-10 | QLI33453 | USA | 2020-05-12 |
| QJR96431 | USA: CA | 2020-03-13 | QNC49349 | Pakistan | 2020-05-15 |
| QKU54102 | USA: Washington,King County | 2020-03-15 | QMT94417 | USA: Washington,Yakima County | 2020-05-27 |
| QLA48060 | USA: NY | 2020-03-24 | QMT54534 | USA: Washington,Yakima County | 2020-06-17 |
| QNG41574 | USA: Minnesota | 2020-03-25 | QLG76514 | Australia: Victoria | 2020-06-20 |
| QKV08176 | USA: Washington,King County | 2020-03-26 | QNG42985 | USA: FL | 2020-06-23 |
| QKV37245 | Australia: Northern Territory | 2020-03-27 | QMT97141 | USA: FL | 2020-06-30 |
| QNI23218 | USA: Virginia | 2020-04 | QNB17780 | Bangladesh | 2020-07-07 |
| QLG99793 | USA: CA | 2020-04-16 | QMU93213 | USA: Wisconsin, Dane county | 2020-07-13 |
| QLY88596 | USA: GA | 2020-04-27 | QNA70543 | Bangladesh | 2020-07-19 |

From Table 1, it was established that the majority of the diversified mutations are deleterious and cause the stability of the protein to decrease, thus indicating the amplification of intricate virulence of SARS-CoV-2.

### Sequence Homology and Mutations of SARS-CoV-2 ORF10

It was reported that SARS-CoV-2 ORF10 is not homologous with other proteins in the NCBI depository^9^. The SARS-CoV-2 ORF10 was blasted in the NCBI depository and no significant homology was detected for ORF10 SARS-CoV as well as Bat-CoV ORF10. Surprisingly, SARS-CoV-2 ORF10 showed 97.37% homology to Pangolin-CoV ORF10 (QIG55954.1 (*Release date: 2020-05-18; Collection date: 2019-03-29; Geo-location: China; Host: Sunda pangolin (Manis javanica))*) (Figure 2)^19^.

Only the serine (S) has been mutated to asparagine (N) at amino acid position 25 in SARS-CoV-2 ORF10 from the Pangolin-CoV ORF10 and the mutation is deleterious (PROVEAN score −3). Due to this mutation, the stability of the protein structure is predicted to be decreased and consequently the intricate virulence of SARS-CoV-2 will escalate.

Analysis of the per-residue intrinsic disorder predispositions of the ORF10 of SARS-CoV-2 and ORF10 proteins from SARS-CoV and Pangolin-CoV provide further evidence of their differences. Figure 3A represents the results of this analysis and shows that while ORF10 proteins from SARS-CoV-2 and Pangolin-CoV show very similar disorder profiles, the per-residue disorder propensity of the ORF10 protein from SARS-CoV is remarkably different, especially within the C-terminal half of this protein. This is in agreement with the results of other analyses conducted in this study.

**Figure 3.**
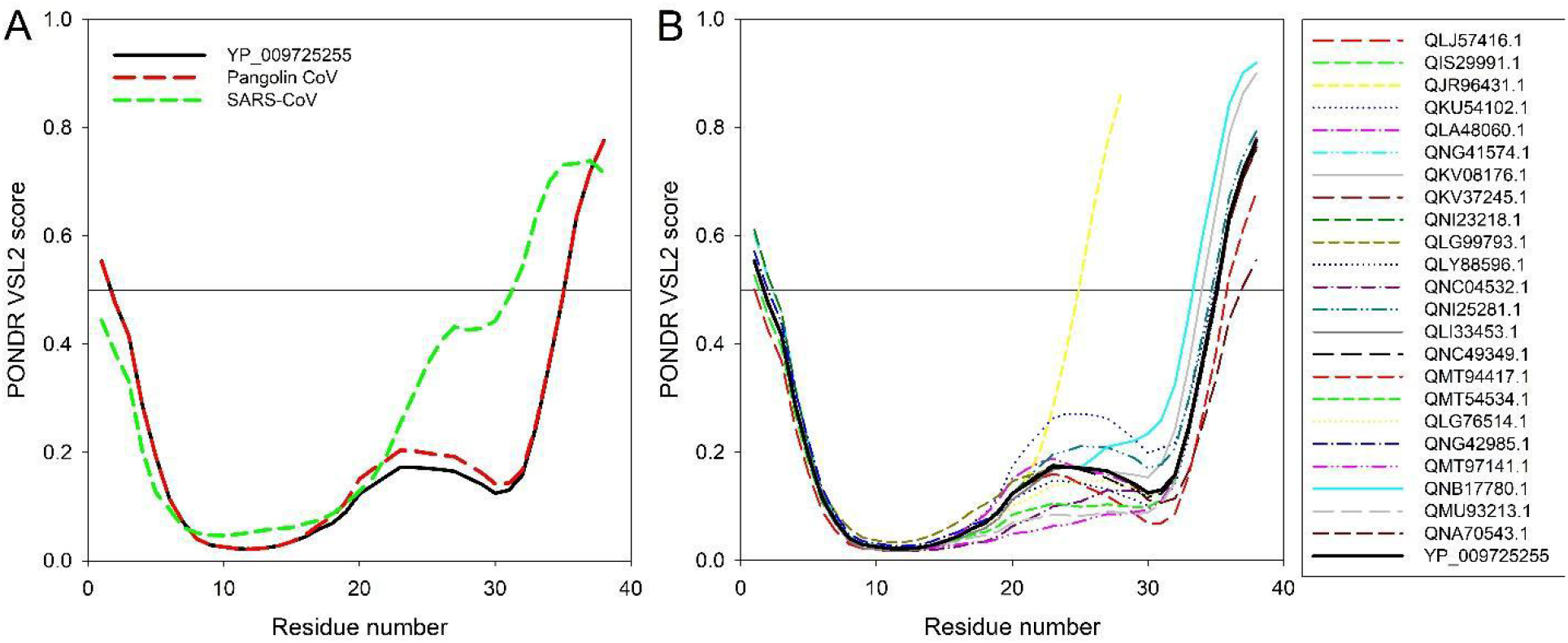
(A) Comparison of the intrinsic disorder profile of the reference ORF10 protein from SARS-CoV-2 (YP_009725255) from the NC_045512 SARS-CoV2 genome (China, Wuhan) (bold black curve) with those of ORF10 proteins from the Pangolin-CoV (QIG55954.1) and SARS-CoV TW-HP1 (UniProt ID: Q6SRY8). (B) Analysis of the intrinsic disorder predisposition of the unique variants of SARS-CoV2 ORF10 in comparison with the reference ORF10 protein from SARS-CoV-2 (YP_009725255) from the NC_045512 SARS-CoV2 genome (China, Wuhan) (bold black curve). Analysis is conducted using PONDR-VSL2 algorithm^23^, which is one of the more accurate standalone disorder predictors^24–26^. A disorder threshold is indicated as a thin line (at score = 0.5). Residues/regions with the disorder scores > 0.5 are considered as disordered, whereas residues with disorder scores between 0.25 and 0.5 are considered highly flexible, and residues with disorder scores between 0.1 and 0.25 are taken as moderately flexible.

Figure 3B compares intrinsic disorder predispositions of the 24 unique variants of ORF10 protein from different isolates of SARS-CoV-2. It is seen that intrinsic disorder predispositions can vary significantly, especially within the C-terminal half of the protein. In fact, majority of substitutions found within the N-terminal region (residues 1-15; i.e., mutations G2D, I4L, V6I, A8V, P10S, I13M, and Y14C) have very little effect on the local intrinsic disorder predisposition of ORF10. On the other hand, ORF10 variants with the mutations within the C-terminal region (residues 20-38; i.e., mutations R20I/K, N22T, S23F, R24C/L, Y26H, A28V, V30A/L, D31Y, F35S, L37P/F, and T38I, as well as shortened QJR96431.1 variant, which is truncated due to a nonsense mutation at the position 29) typically show rather substantial variability in their local disorder predispositions. The most significant changes are observed within the “disorder hump” region (residues 20-30), intensity of which is increased in QKU54102.1 (Y26H), QNI25281.1 (V30A), and QNB17780.1 (F35S) ORF10 variants, whereas in the variants QMT54534.1 (R24L), QNC04532.1 (R20I), QMU93213.1 (R24C), and QMT97141.1 (S23F), this hump is either eliminated or noticeably flattened. Interestingly, comparison of the Figure 3A and 3B shows that the variability in the disorder predisposition between many variants of the ORF10 protein from various SARS-CoV-2 isolates is noticeably greater than that between the reference ORF10 from SARS-CoV-2 and ORF10 from Pangolin-CoV. On the other hand, none of the SARS-CoV-2 ORF10 variants (with the exception for the truncated QJR96431.1 variant) has as disordered C-terminal half as the ORF10 protein from SARS-CoV does.

### Comparison of SARS-CoV2 ORF10 and Pangolin-CoV ORF10

Considering the highest amount of sequence homology of ORF10 proteins of SARS-CoV-2 and Pangolin-CoV, we intended to discover the parity and disparity between the ORF10 proteins of SARS-CoV-2 and Pangolin-CoV. We, therefore, performed a multi-dimensional analysis of both ORF10 proteins from structural, physicochemical, biophysical and immunological aspects to understand the origin of SARS-CoV-2 from the ORF10 perspective.

Exploration of similarities between SARS-CoV-2 and Pangolin ORF10 sequences (Figure 4A) revealed that neither of them had disulfide linkages. However, many differences were detected. The SARS-CoV-2 ORF10 protein was classified as an alpha-helical transmembrane protein (with probability 0.489) owing to the server ABTMpro as well as the presence of a majority of hydrophobic amino acids, whereas the Pangolin-CoV ORF10 sequence was predicted to be a non-transmembrane protein (with probability 0.513). Also, it was discerned that the predicted probability of antigenicity of SARS-CoV-2 ORF10 was slightly higher than that of Pangolin-CoV ORF10. It was predicted that both proteins are located in the capsid region of the virus as both of them have a positive distance score, with a higher score for Pangolin-CoV (0.1502) than for SARS-CoV-2 (0.1141).

**Figure 4.**
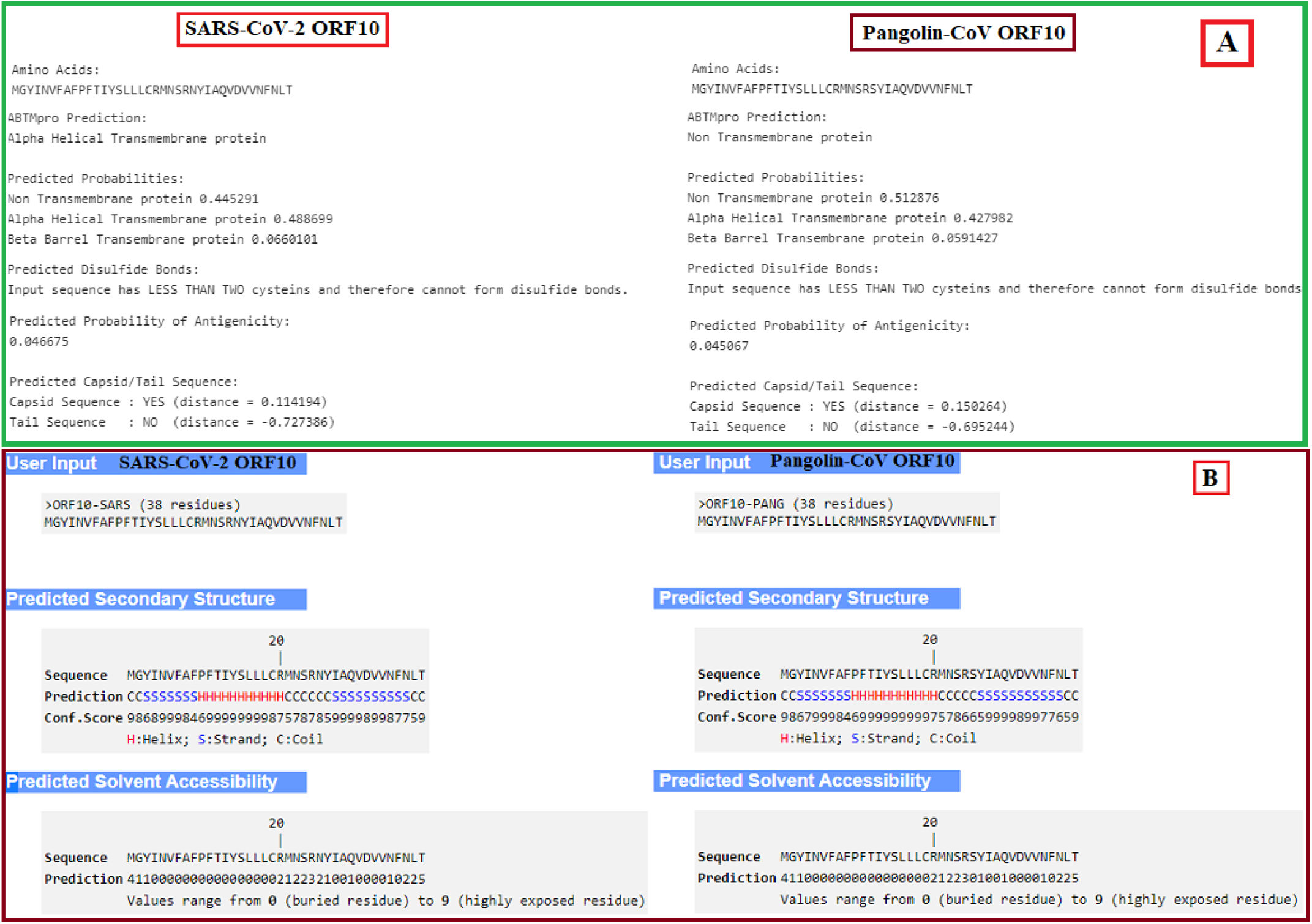
(A): Basic properties of ORF10 proteins of SARS-CoV-2 and Pangolin-CoV; (B): Peptide and solvent accessibility properties of ORF10 proteins of SARS-CoV-2 and Pangolin-CoV

To achieve deeper insights into the ORF10 proteins of SARS-CoV-2 and Pangolin-CoV, we characterized their secondary structure (Figure 4B) and found them to be very much similar except for a significant difference at the position 26, Tyr (Y), which for SARS-CoV-2 ORF10 is in the coil region whereas for Pangolin-CoV ORF10, it is located in the strand region. Most of the residues, 23 in SARS-CoV-2 ORF10 and 24 in Pangolin-CoV ORF10, are buried and consequently, the solubility of SARS-CoV-2 ORF10 is slightly higher than that of Pangolin-CoV.

After structural and fundamental property studies, a subsequent thorough analysis of the physicochemical properties of two ORF10 proteins of SARS-CoV-2 and Pangolin-CoV was performed, which unveiled the high similarity based on extinction coefficient, isoelectric point and net charge (Figure 5A). However, the molecular weight (4449.18 g/mol) of the SARS-CoV-2 ORF10 was higher compared to Pangolin-CoV ORF10 (4422.16 g/mol), due to the substitution of S (low molecular weight) of Pangolin-CoV to N (high molecular weight) of SARS-CoV-2. The enzyme cleavage sites for the SARS-CoV-2 and Pangolin-CoV ORF10 were also indistinguishable for all proteases (Figure 5B).

**Figure 5.**
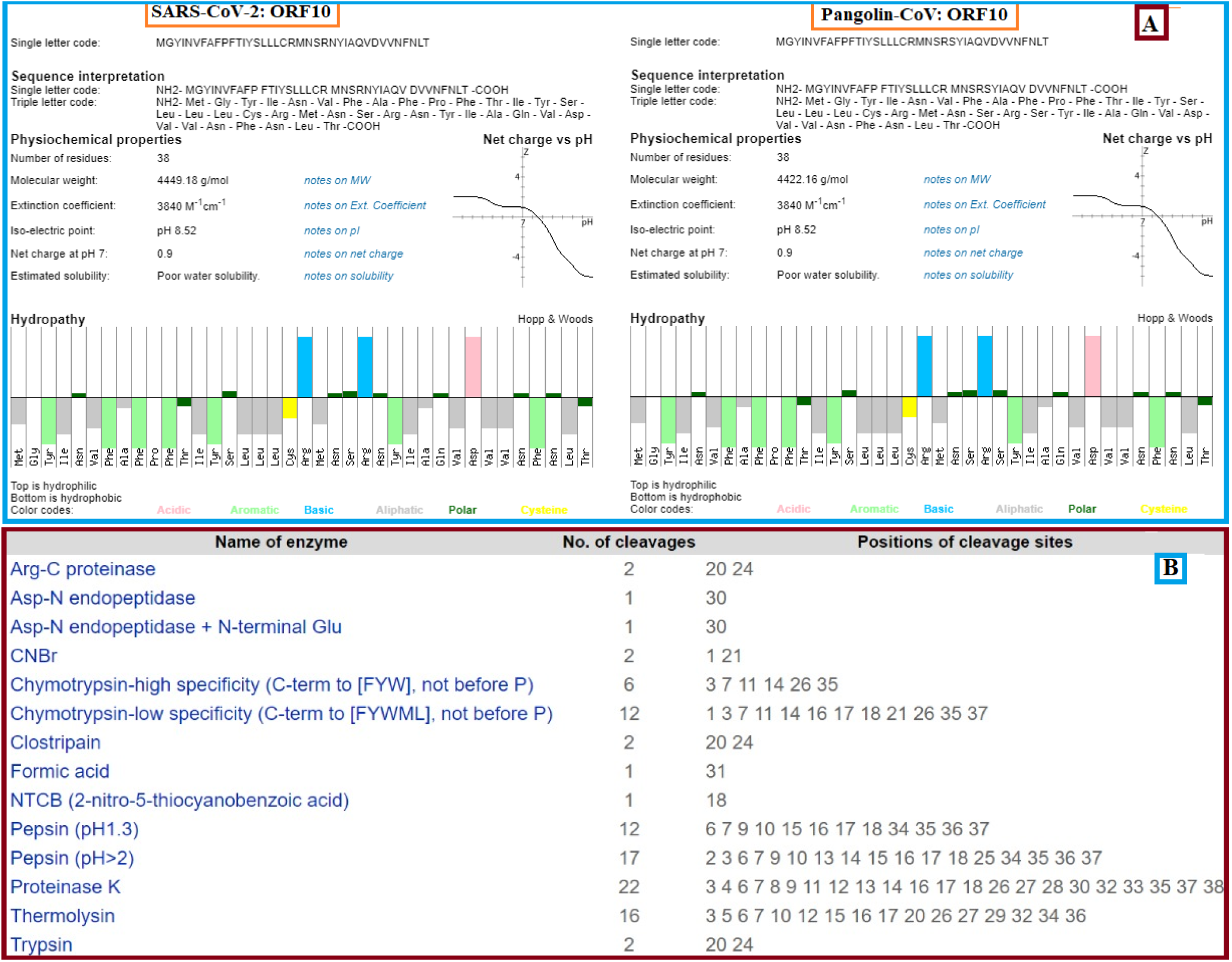
(A): Physicochemical properties and hydropathy of ORF10 of SARS-CoV-2 and Pangolin-CoV; (B): Enzymes and numbers of associated cleavages and their positions.

Protein intrinsic disorder analysis disclosed the presence of hotloops in both sequences within the same span of amino-acids (26-38). However, the presence of loops/coils (22-29) was a distinct characteristic of SARS-CoV-2 ORF10 and no such structures were observed for Pangolin ORF10 (Figure 6).

**Figure 6.**
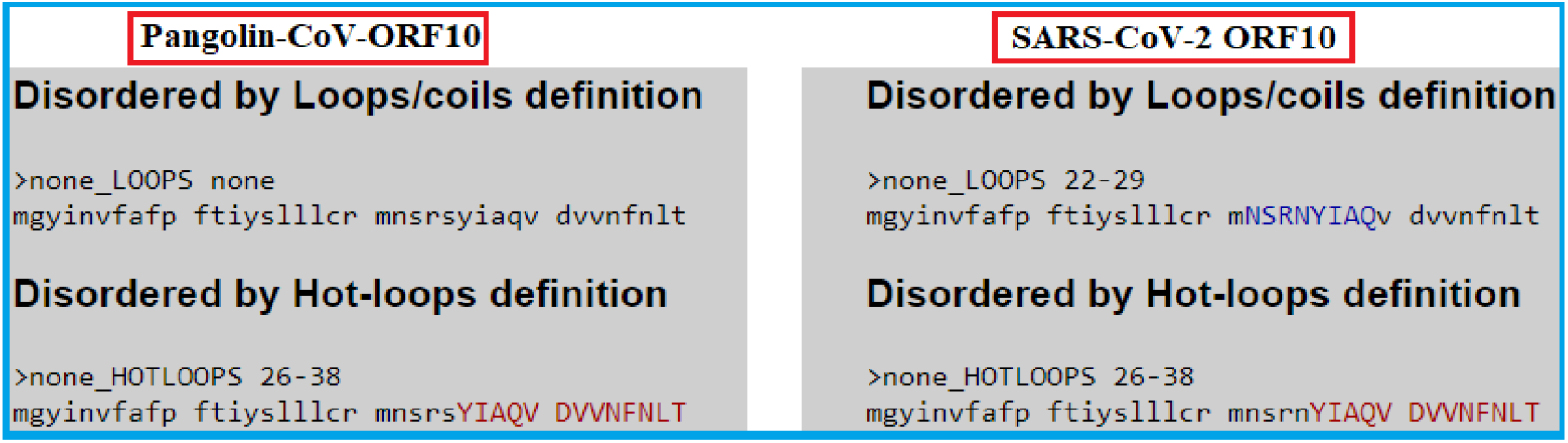
Disordered loops and hotloops of ORF10 of SARS-CoV-2 and Pangolin-CoV

To shed light on the immunogenic properties of ORF10, we carried out immunoinformatics analysis and identified (Figure 7) nine amino acid long epitopes in 11 Cytotoxic T-lymphocytes (CTLs) from the SARS-CoV-2 ORF10 sequence across all 12 HLA subtypes. Their scores were recorded, and corresponding epitope-bearing mutations were analysed. Comparison of scores with the original epitopes were done and thereby predicted the increase/decrease in binding affinity for class I MHC molecules due to mutations. These eleven epitopes and mutational sequence-bearing epitopes were analysed using the IDEB tool to account for their immunogenicity.

**Figure 7.**
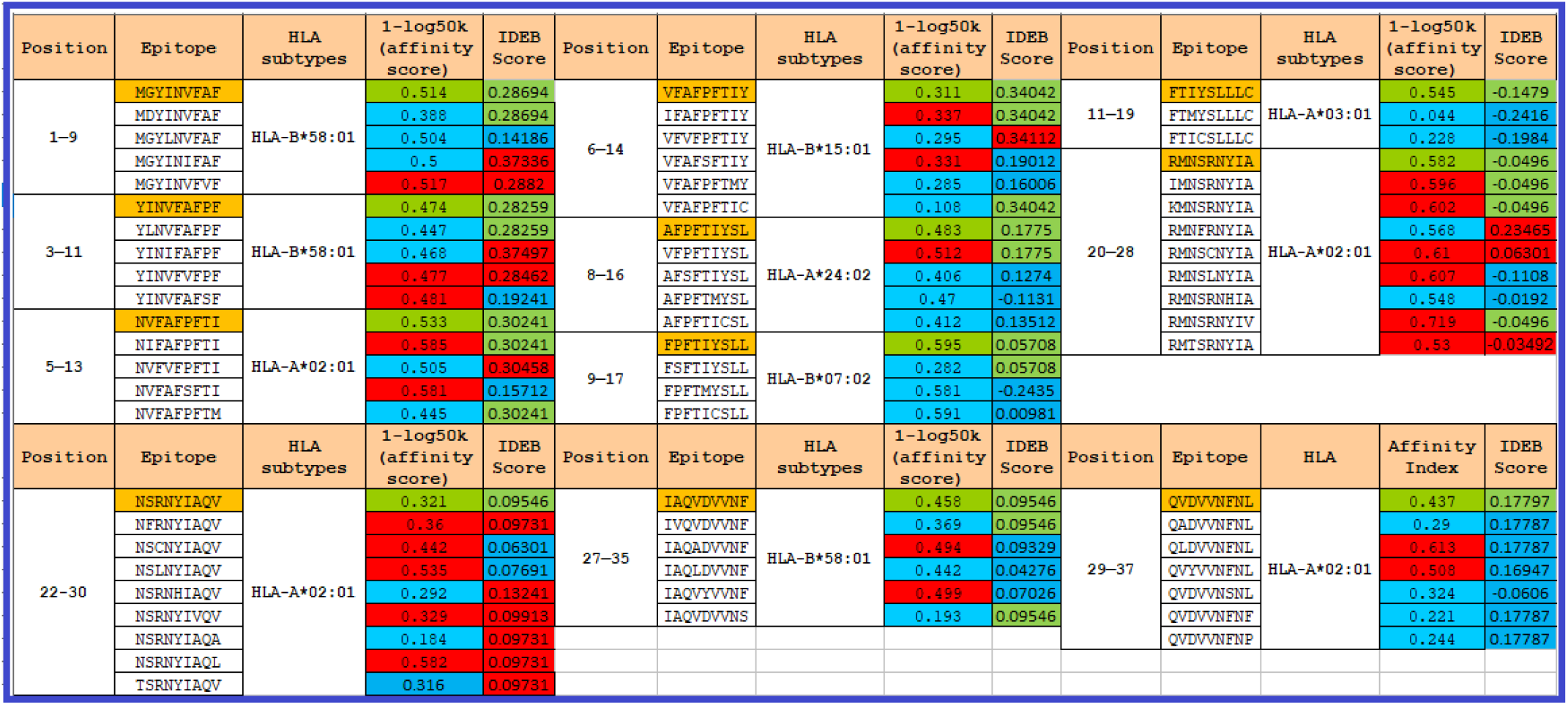
Eleven distinct epitopes in the SARS-CoV-2 ORF10 were identified and analysed for binding affinity using PICKPOCKET across 12 HLA subtypes. The IDEB score was predicted using the IDEB immunogenicity tool. Eleven epitopes (marked in orange) from the Wuhan SARS-CoV-2 ORF10 sequence. Scores in red/blue show an increase/decrease concerning the score associated with nine epitopes. Green marked scores convey the immunogenicity value remaining unchanged.

## Discussion

A detailed study of the ORF10 protein was carried out to evaluate its potential to yield to variants that could possibly alter viral pathogenicity. It was observed that each SARS-CoV-2 ORF10 sequence possesses one distinct mutation. Each of the twenty-two SARS-CoV-2 ORF10 variants is at a uniquely different position. None of these mutations in the SARS-CoV-2 ORF10, however, contributes to the determination of clades of SARS-CoV-2. Of all variants, a total of 13 variants were identified to possess mutations at amino acid positions 22-38 and in a region predicted to contain overlapping loops/coils and hot-loop regions of the ORF10 protein. All mutations were predicted to be deleterious with decreased effect on protein structure stability except S23F, which increased stability, denoting that these mutations play an active role in enhancing intrinsic propensity disorder (IPD) and allowing the protein to undergo more favorable interactions with other proteins. Two other mutations, I4L and V6I, were found to be in the MoRF region of ORF10, and which may also possibly contribute to the IPD as well.

The mutations at positions 20 and 24 were also significant due to their sensitivity for trypsin activity. Four ORF10 variants (QNC04532.1, QMT54534.1, QMU93213.1 and QLA48060.1) possess four mutations at these two positions. Among them, three variants harboring the mutations R20I, R24L and R24C provide trypsin resistance, while the fourth variant (QLA48060.1) with the R20K mutation is susceptible to protease degradation.

An amino acid homology of 97.37% was observed between SARS-CoV-2 ORF10 and Pangolin-CoV ORF10. Although most physicochemical and peptide properties are similar, the probability of antigenicity is greater for SARS-CoV-2 ORF10 than that of Pangolin-CoV ORF10 and consequently a stronger immune response is predicted for SARS-CoV-2 ORF10. A change from strand (Pangolin-CoV ORF10) to coil (SARS-CoV-2 ORF10) at position 26 (tyrosine (Y)), is predicted indicating the higher disordered state of the protein. A sequence with the Y26H mutation was also detected in SARS-CoV-2 ORF10, which showed that a hydrophobic amino acid was replaced by a hydrophilic amino acid, thus increasing the probability for more ionic interactions.

Analysis identified ORF10 mutations predicted to alter binding affinity to respective HLA alleles and to possibly correspondingly change the immunogenicity of SARS-CoV-2 ORF10. Eight ORF10 variants (containing one of the following mutations each G2D, I4L, I13M, Y14C, Y26H, F35S, L37S and L37P (Table 1)) accounted for 40% of total mutations and demonstrated decreased affinity for MHC class I, 25% of the variants (carrying mutations R20K, R20I, R24C, R24L and D31Y) predict for increased affinity, and 35% of the variants (carrying mutations V6I, A8V, P10S, S23F, A28V and V30A) contain both high and low binding affinity epitopes. This may indicates that mutations in ORF10 are predominantly decrease the affinity of epitopes to escape the host-immune system, while in the mixed cases the effect of increased affinity by mutations is nullified by the presence of mutations contributing to decreased affinity. For mutations showing only increased binding affinity epitopes, it is hypothesized that acquiring more than one mutation in a single sequence in the future will nullify them as well. In addition, the immunogenicity score prediction revealed that a large number of mutations had decreased or no effect and very few of them exhibited an increased immunogenicity score, which may be a possible strategy adopted by SARS-CoV-2 to evade the host-immune response. Six mutation-bearing sequences (QLJ57416.1, QMT97141.1, QLY88596.1, QNC49349.1, QMT54534.1, and QLG76514.1) were found to contain epitopes showing both high affinity binding for MHC class 1 and high immunogenicity, indicating that these epitopes can mount significant immune response and might serve as potential targets for vaccine candidates. More critical study in ORF10 SARS-CoV2 is necessary to monitor high frequency mutations that could change viral pathogenesis.

ORF10 protein of SARS-CoV-2 and Pangolin-CoV are similar. However, there are predicted notable differences detected between these two ORF10 proteins in terms of loop/coil structure, antigenicity, solubility, and in mutational diversification of SARS-CoV-2. These significant disagreements of various physicochemical, structural, immunological properties despite an amino acid homology (97.37%) between the ORF10 proteins of SARS-CoV-2 and Pangolin-CoV are quite surprising, and deserving of further study.

## Data and Methods

### Data acquisition

There were 11,288 complete genomes of SARS-CoV-2 available on the NCBI (National Center for Biotechnology Information) database, as of 28th August 2020. Each genome contains the ORF10 accessory protein and among them only 34 sequences were found to be unique. Among these unique ORF10 protein sequences, only 22 sequences possess only one missense mutation each and the remaining sequences possess ambiguous mutations. It is noted that, there was only one ORF10 sequence (QJR96431.1) which was truncated due to a nonsense mutation at amino acid position 29. The present study focused on these 23 ORF10 proteins (Table 2).

A reference ORF10 protein (YP_009725255.1) of the SARS-CoV-2 genome (NC_045512) from Wuhan, China was used to identify the mutations^27^.

The miscellany of ORF10 variants of SARS-CoV-2 is clearly observed in the sequence-based homology (Figure 8(A)) and phylogeny (Figure 8(B)).

**Figure 8.**
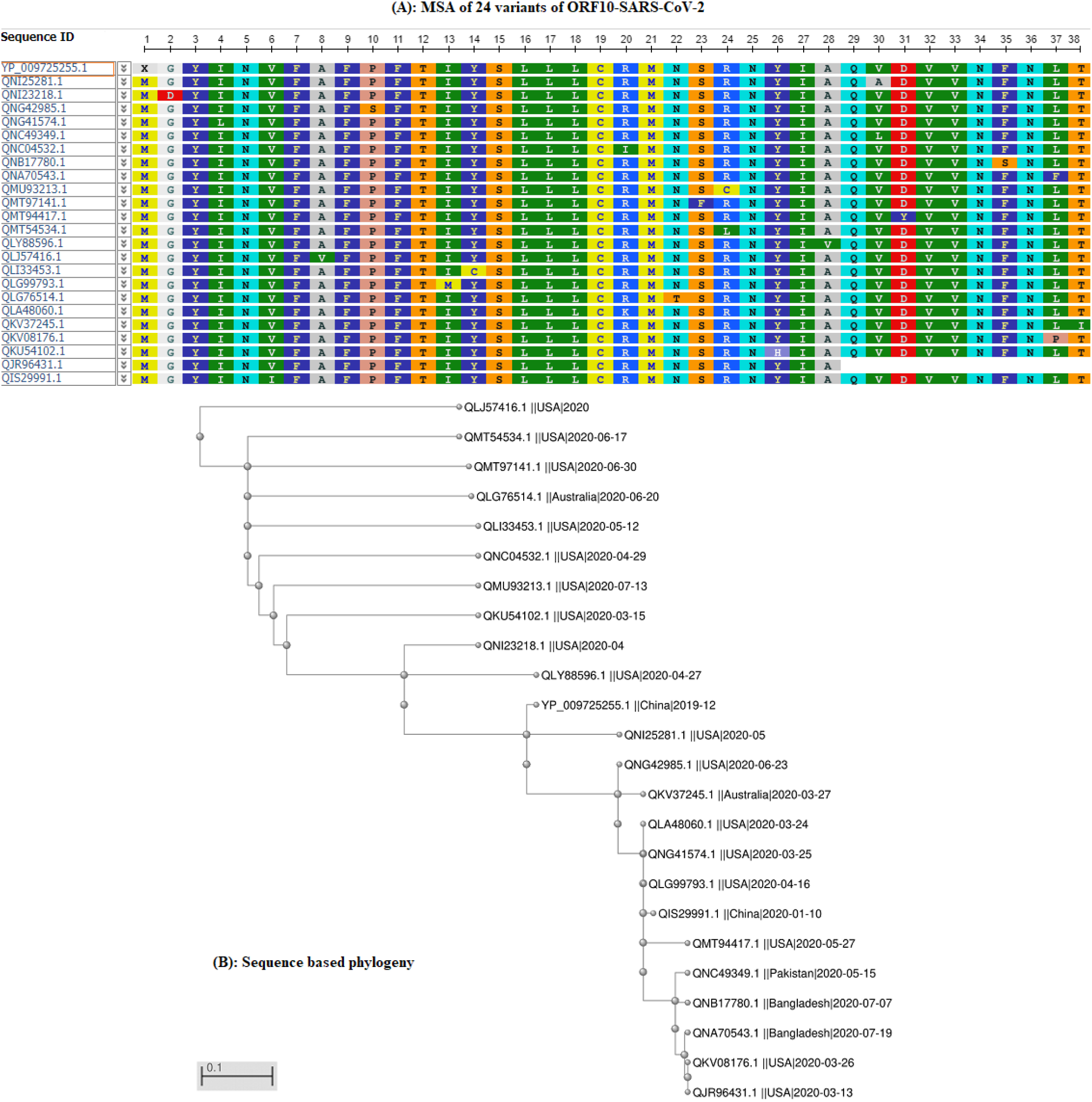
(A): Multiple sequence alignment (MSA) of 24 SARS-CoV-2 ORF10 proteins; (B): Phylogeny of 24 SARS-CoV-2 ORF10 sequences.

Each ORF10 of SARS-CoV-2 is different from the Wuhan SARS-CoV-2 ORF10 sequence utilizing a single amino acid change at a distinct position. Noticeably, these positions (18) are widely varying from the position 2 to 38 for the 22 SARS-CoV-2 ORF10 variants.

## Methods

### Webserver based predictions

The prediction of various properties of ORF10 proteins was determined by several webservers which are briefly described as follows.

- For the prediction of the effect of identified mutations, the PROVEAN webserver was used and also for the structural effects of mutations, and another webserver, I-MUTANT, was used^28–30^. The QUARK webserver was used for the prediction of secondary structure of ORF10 proteins^31–33^.
- Given an amino acid sequence, the ABTMpro webserver predicts whether the given sequence is a transmembrane protein. If the given sequence is a transmembrane protein, it further predicts the probabilities of the protein being an alpha-helix transmembrane protein or a Beta Barrel transmembrane protein. In addition, for various peptide property findings, the INNOVAGEN webserver was used^34^.
- The DIpro can predict whether the given protein sequence contains a cysteine disulfide bond, based on 2D recurrent neural network, support vector machine, graph matching and regression algorithms^35^.
- The protein antigenicity is predicted using the webserver ANTIGENpro, which is a sequence-based, alignment-free and pathogen-independent predictor. A two-stage architecture makes the probability of prediction based on multiple representations of the primary sequence and five machine learning algorithms^36^. The intrinsic disorder prediction of a given protein sequence was made using the server DisEMBL^37^.
- Epitopes of a given amino acid sequence were spotted and analyzed for binding affinity using across 12 HLA (human leukocyte antigen) subtypes (HLA-A*01:01, HLA-A*02:01, HLA-A*03:01, HLA-A*24:02, HLA-A*26:01, HLA-B*07:02, HLA-B*08:01, HLA-B*27:05 B*39:01, B*40:01, B*58:01 and B*15:01). The IDEB (The Immune Epitope Database) score was predicted using the IDEB immunogenicity tool^38,39^.

### Evaluating the per-residue predisposition of various ORF10 proteins for intrinsic disorder

Per-residue disorder distribution within ORF10 protein sequences was evaluated by *PONDR* − *VSL*2^23^, which is one of the more accurate standalone disorder predictors^24–26^. The per-residue disorder predisposition scores are on a scale from 0 to 1, where values of 0 indicate fully ordered residues, and values of 1 indicate fully disordered residues. Values above the threshold of 0.5 are considered disordered residues, whereas residues with disorder scores between 0.25 and 0.5 are considered highly flexible, and residues with disorder scores between 0.1 and 0.25 are taken as moderately flexible.

## Acknowledgments

Authors acknowledge the NCBI sequence (SARS-CoV-2 and Pangolin-CoV-2020) depositors.

## Author contributions statement

SSH conceived the problem and experiment(s). DA, SG, SSH, VNU examined the mutations. SSH, PPC, DA, SG and VNU analysed the results. SSH wrote the primary draft of the article. All authors reviewed, edited and approved the final manuscript.

## References

1. Kong, W.-H. et al. Sars-cov-2 detection in patients with influenza-like illness. Nat. microbiology 5, 675–678 (2020).

2. Ju, B. et al. Human neutralizing antibodies elicited by sars-cov-2 infection. Nature 584, 115–119 (2020).

3. Yousefzadegan, S. & Rezaei, N. Case report: Death due to covid-19 in three brothers. The Am. J. Trop. Medicine Hyg. 102, 1203–1204 (2020).

4. Xu, Z. et al. Pathological findings of covid-19 associated with acute respiratory distress syndrome. The Lancet respiratory medicine 8, 420–422 (2020).

5. Li, G. et al. Coronavirus infections and immune responses. J. medical virology 92, 424–432 (2020).

6. Corman, V. M. et al. Rooting the phylogenetic tree of middle east respiratory syndrome coronavirus by characterization of a conspecific virus from an african bat. J. virology 88, 11297–11303 (2014).

7. Hui, D. S. et al. The continuing 2019-ncov epidemic threat of novel coronaviruses to global health—the latest 2019 novel coronavirus outbreak in wuhan, china. Int. J. Infect. Dis. 91, 264–266 (2020).

8. of the International, C. S. G. et al. The species severe acute respiratory syndrome-related coronavirus: classifying 2019-ncov and naming it sars-cov-2. Nat. Microbiol. 5, 536 (2020).

9. Koyama, T., Platt, D. & Parida, L. Variant analysis of sars-cov-2 genomes. Bull. World Heal. Organ. 98, 495 (2020).

10. Kiyotani, K., Toyoshima, Y., Nemoto, K. & Nakamura, Y. Bioinformatic prediction of potential t cell epitopes for sars-cov-2. J. Hum. Genet. 65, 569–575 (2020).

11. Giri, R. et al. Understanding covid-19 via comparative analysis of dark proteomes of sars-cov-2, human sars and bat sars-like coronaviruses. Cell. Mol. Life Sci. 1–34 (2020).

12. Uversky, V. N. Multitude of binding modes attainable by intrinsically disordered proteins: a portrait gallery of disorder-based complexes. Chem. Soc. Rev. 40, 1623–1634 (2011).

13. Gordon, D. E. et al. A sars-cov-2 protein interaction map reveals targets for drug repurposing. Nature 1–13 (2020).

14. Díaz, J. Sars-cov-2 molecular network structure. Front. Physiol. 11, 870 (2020).

15. Liang, Q. et al. Virus-host interactome and proteomic survey of pmbcs from covid-19 patients reveal potential virulence factors influencing sars-cov-2 pathogenesis. bioRxiv (2020).

16. Cagliani, R., Forni, D., Clerici, M. & Sironi, M. Coding potential and sequence conservation of sars-cov-2 and related animal viruses. Infect. Genet. Evol. 104353 (2020).

17. Le Bert, N. et al. Sars-cov-2-specific t cell immunity in cases of covid-19 and sars, and uninfected controls. Nature 584, 457–462 (2020).

18. Kim, D. et al. The architecture of sars-cov-2 transcriptome. Cell (2020).

19. Liu, P. et al. Are pangolins the intermediate host of the 2019 novel coronavirus (sars-cov-2)? PLoS Pathog. 16, e1008421 (2020).

20. Johnson, M. et al. Ncbi blast: a better web interface. Nucleic acids research 36, W5–W9 (2008).

21. Madeira, F. et al. The embl-ebi search and sequence analysis tools apis in 2019. Nucleic acids research 47, W636–W641 (2019).

22. Malhis, N., Jacobson, M. & Gsponer, J. Morfchibi system: software tools for the identification of morfs in protein sequences. Nucleic acids research 44, W488–W493 (2016).

23. Obradovic, Z., Peng, K., Vucetic, S., Radivojac, P. & Dunker, A. K. Exploiting heterogeneous sequence properties improves prediction of protein disorder. Proteins: Struct. Funct. Bioinforma. 61, 176–182 (2005).

24. Meng, F., Uversky, V. N. & Kurgan, L. Comprehensive review of methods for prediction of intrinsic disorder and its molecular functions. Cell. Mol. Life Sci. 74, 3069–3090 (2017).

25. Peng, Z.-L. & Kurgan, L. Comprehensive comparative assessment of in-silico predictors of disordered regions. Curr. Protein Pept. Sci. 13, 6–18 (2012).

26. Fan, X. & Kurgan, L. Accurate prediction of disorder in protein chains with a comprehensive and empirically designed consensus. J. Biomol. Struct. Dyn. 32, 448–464 (2014).

27. Wu, X. et al. Co-infection with sars-cov-2 and influenza a virus in patient with pneumonia, china. Emerg. infectious diseases 26, 1324 (2020).

28. Choi, Y., Sims, G. E., Murphy, S., Miller, J. R. & Chan, A. P. Predicting the functional effect of amino acid substitutions and indels. PloS one 7, e46688 (2012).

29. Choi, Y. A fast computation of pairwise sequence alignment scores between a protein and a set of single-locus variants of another protein. In Proceedings of the ACM Conference on Bioinformatics, Computational Biology and Biomedicine, 414–417 (2012).

30. Choi, Y. & Chan, A. P. Provean web server: a tool to predict the functional effect of amino acid substitutions and indels. Bioinformatics 31, 2745–2747 (2015).

31. Xu, D. & Zhang, Y. Ab initio protein structure assembly using continuous structure fragments and optimized knowledge-based force field. Proteins: Struct. Funct. Bioinforma. 80, 1715–1735 (2012).

32. Xu, D. & Zhang, Y. Toward optimal fragment generations for ab initio protein structure assembly. Proteins: Struct. Funct. Bioinforma. 81, 229–239 (2013).

33. Hassan, S. S., Choudhury, P. P., Basu, P. & Jana, S. S. Molecular conservation and differential mutation on orf3a gene in indian sars-cov2 genomes. Genomics 112, 3226–3237 (2020).

34. Cheng, J., Randall, A. Z., Sweredoski, M. J. & Baldi, P. Scratch: a protein structure and structural feature prediction server. Nucleic acids research 33, W72–W76 (2005).

35. Cheng, J., Saigo, H. & Baldi, P. Large-scale prediction of disulphide bridges using kernel methods, two-dimensional recursive neural networks, and weighted graph matching. Proteins: Struct. Funct. Bioinforma. 62, 617–629 (2006).

36. Magnan, C. N. et al. High-throughput prediction of protein antigenicity using protein microarray data. Bioinformatics 26, 2936–2943 (2010).

37. Linding, R. et al. Protein disorder prediction: implications for structural proteomics. Structure 11, 1453–1459 (2003).

38. Zhang, H., Lund, O. & Nielsen, M. The pickpocket method for predicting binding specificities for receptors based on receptor pocket similarities: application to mhc-peptide binding. Bioinformatics 25, 1293–1299 (2009).

39. Vita, R. et al. The immune epitope database (iedb): 2018 update. Nucleic acids research 47, D339–D343 (2019).

